# Self-Supervised Missing Wedge Correction in Soft X-Ray Tomography: Towards Accurate Cellular Morphology and Volume Quantification

**DOI:** 10.64898/2025.12.23.696166

**Authors:** ShaoSen Chueh, Mary Lopez-Perez, Charlotte de Ceuninck van Capelle, Takashi Ishikawa, Alessandro Zannotti, Victoria Castro, Gema Calvo, Pablo Gastaminza, Conal McIntyre, David Rogers, Stephen O’Connor, Martina Donnellan, Jeremy C. Simpson, Sergey Kapishnikov

**Author notes:** Correspondence: ShaoSen Chueh and Sergey Kapishnikov.

## Abstract

Soft X-Ray tomography (SXT) is a non-invasive bio-imaging technique that enables 3D imaging of cellular structures in large volume, with a unique resolution range that bridges the gap between fluorescence and transmission electron microscopy. However, a fundamental limitation, the missing wedge artefact caused by incomplete tilt-series acquisition, introduces systematic structural elongation in the reconstructed tomograms. This artifact compromises accurate quantitative biological analysis by overestimating cellular and organelle volumes. To overcome this persistent issue, we introduce a novel, self-supervised missing wedge correction model that learns key sine-wave patterns from existing SXT sinograms of the tilt series stacks. This model can be applied to recover the missing-angle region of the sinogram, reducing distortions and elongations in the reconstructed tomograms. We demonstrate a significant quantitative improvement in artifact removal, achieving faithful recovery of the spherical morphology of lipid droplets. We further applied this method to *Plasmodium falciparum* hemozoin crystals, a biomarker in antimalarial drug efficacy studies. Our model successfully reduced volume overestimation, achieving up to a 16% decrease in distorted volume. This level of precision is paramount for correctly interpreting the mode of action of antimalarial drugs.

## Introduction

Soft X-Ray Tomography (SXT) is a non-invasive imaging technique that enables high-resolution label-free imaging of cells and tissues in their near-native states. Its ability to visualize three-dimensional cellular ultrastructures allows researchers to quantify cellular volumes and morphological features important in biological and biomedical research [1][2][3][4][5][6][7]. One significant application of SXT is in studying the mode of action of antimalarial drugs that inhibit hemozoin crystal growth, a process used by malaria parasites to detoxicate heme [8][9][10]. Quantifying the volume and morphology of hemozoin crystals imaged by SXT has been key to exploring the role of antimalarial drugs, such as quinolines, in inhibiting the growth of hemozoin crystals, and how this effect becomes lethal to malaria parasites [11][12][13][14][15][16][17][18][19].

The vast majority of biological samples imaged by SXT are supported on flat specimen carriers, such as standard electron microscopy grids. However, SXT imaging of flat samples is affected by the missing wedge effect, which arises from the blocking of X-ray transmission at high tilt angles by the sample carrier or the sample itself. This constitutes an impediment to accurate quantification of cellular structures in SXT, and across various microscopy imaging modalities [20][21][22][23][24][25][26]. The achievable sample tilt range is determined by the physical constraints of the specimen holders and is typically ±60° to ±70°, or in rare cases greater, depending on the instrument [27][28][29]. Consequently, the information at the sample tilts outside that range is missing and cannot be attained, resulting in elongation artifacts and vanishing contrast along the projection direction in the tomogram reconstructed from tilt series [30][31][32]. The presence of elongation artifacts will lead to falsely enlarged cellular volumes and altered morphology due to the loss of information. Accurate quantification of cellular volume and shape is crucial for addressing biological and biomedical questions through imaging, such as understanding the mode of action of antimalarial drugs.

Multiple attempts have been made to mitigate the effects of the missing information in tomographic reconstruction. Algorithmic methods [33][34] are susceptible to introducing artifacts, especially when analyzing complex biological cellular structures. Learning-based mitigation methods also face inherent limitations. A high proportion of proposed models depend on either ground truth data [35][36][37][38] or the assumption of target shape [39][40] for training. These prerequisites constrain their usability in practical bio-imaging scenarios. While approaches such as IsoNet [41] and spIsoNet [42] were developed for single-particle Cryo-Electron Microscopy (Cryo-EM) missing wedge correction without requiring external ground truth, their reliance on rotated subtomograms as training targets renders them more suitable for repeating or locally consistent structures for subtomograms extraction [43][44][45][46]. Furthermore, the requisite subtomogram extraction and iterative refinement processes impose substantial computational overhead. UsiNet [47] introduced a practical self-supervised sinogram inpainting technique for missing wedge correction that eliminates the need for ground-truth data during training, a significant advantage given the typical absence of full-angle tilt series. However, the UsiNet framework fails on the heterogeneous sinogram of SXT. We surmise that the major reason for this is the use of a simple Mean Squared Error (MSE) loss that smooths out complex details, resulting in blurry, non-diagnostic output. Furthermore, the 2-step inpainting of the sinogram is too computationally demanding in terms of both time complexity and GPU memory for practical use in 3-dimensional SXT data.

Therefore, we propose a 1-step self-supervised sinogram inpainting framework based on Generative Adversarial Networks (GAN) [48][49][50][51][52], which addresses the over-smoothing effect of MSE loss [53]. Furthermore, our method also significantly reduces the computational resources needed by training the model to recover only the masked region (30 projection slices, instead of the entire tilt series stack), while leaving the inpainting of the actual missing-wedge region to the inference phase. This modification is based on the fundamental premise of self-supervised sinogram inpainting: the model can effectively learn the underlying sine wave patterns even with only a small portion of the sinogram, and apply this knowledge to recover missing sinogram regions.

We validate our approach using two distinct examples. Firstly, by recovering the spherical shapes of lipid droplets, showing a significant improvement in eccentricity and axis ratio (minor-axis/major-axis), as proof that our method faithfully restored the true shapes of cellular structures. Secondly, we demonstrate our model’s effectiveness with a diverse range of cellular structures, including malaria parasites in red blood cells. Our approach mitigates the missing-wedge elongation and contrast vanishing artifacts in reconstructed tomograms, allowing a more accurate observation of sample morphology and volumetric quantification, including hemozoin crystals. For malaria research, this improvement will enable a better quantification of the growth of hemozoin crystals, an indicator of the effectiveness of antimalarial drugs such as quinolines [11][12][13][14][15][16][17][18][19].

## Results

### Self-Supervised Training of Sinogram Inpainting

In contrast to UsiNet, we simplified the training pipeline by focusing solely on recovering the manually masked region, where the ground truth is available for self-supervised training. The soft X-ray microscopy tilt series typically comprises 110 to 140 projection slices (±55° to ±70° angular range). For this study, we experimented with recovering 40 and 60 angular degrees (20 or 30 degrees from each side of the sinogram). We denoted the number of degrees recovered from a single side of the sinogram as N, i.e., when inpainting 40 degrees overall, the model recovers 20 degrees from each side.

During training, a random 2N-slice subset is extracted from the original tilt series (Fig. 1a). To simulate the missing angular wedge encountered during inference, a strategic masking scheme is employed: either the upper or the lower N slices of the cropped 2N-slice sinogram are masked (i.e., pixel values set to zero). The model is then trained exclusively to recover this masked region. This deliberate masking of an edge region (upper or lower half) is crucial because the missing angles to be recovered during inference are located at the edges of the complete sinogram. Randomly masking an internal 30-slice region would allow the model to learn recovery cues from both sides of the gap, which is not representative of the inference scenario where only one side of the missing angle boundary provides contextual information.

**Figure 1.**
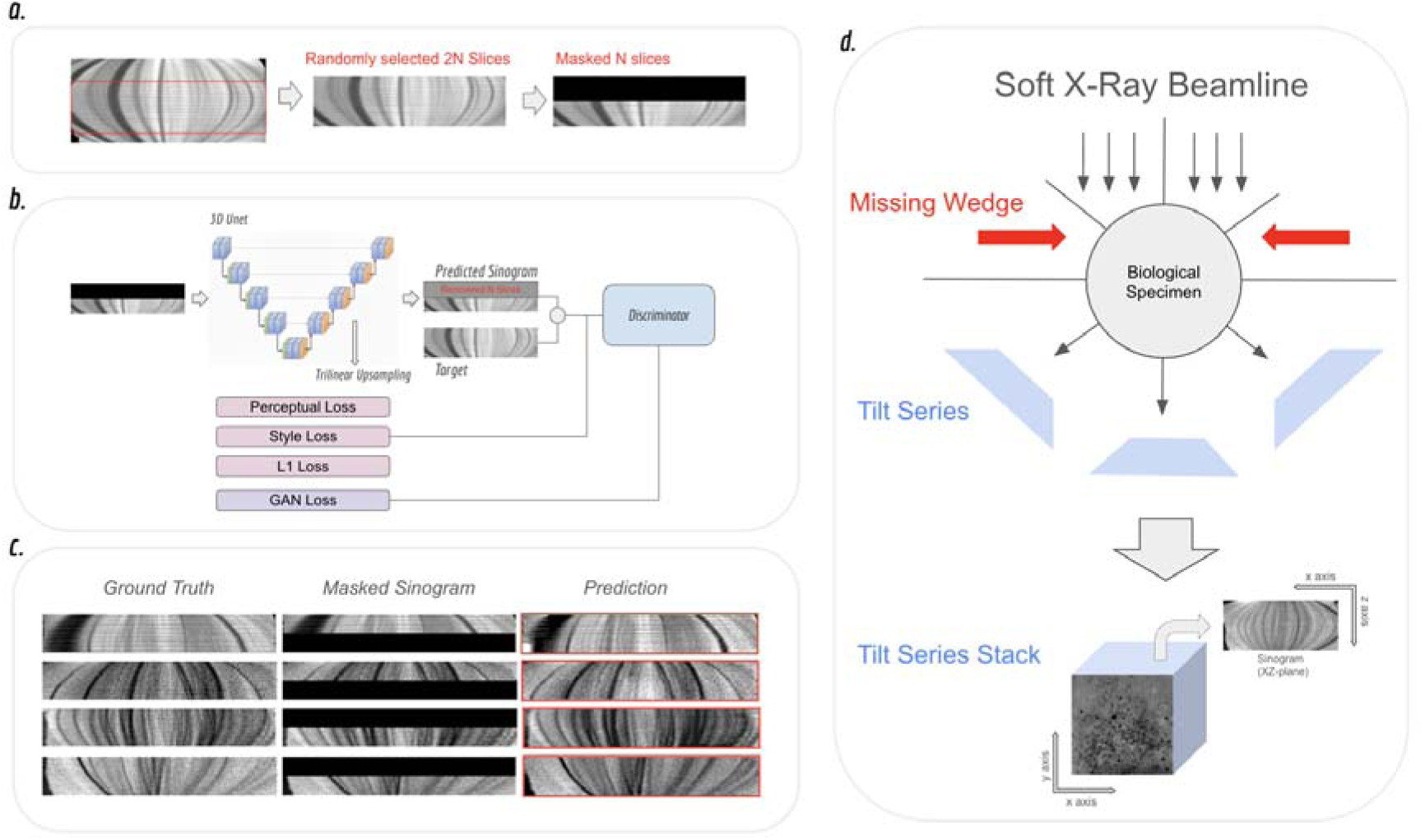
The self-supervised sinogram inpainting training pipeline. a) To prepare the training data, a random 2N-slice subset is first cropped from the available tilt series stack (which usually spans ±55 to ±60 degrees). Then, a manual mask is applied to either the upper or lower half of this cropped sinogram (i.e., N slices) to simulate the missing angular wedge boundary. This masked region serves as the ground truth for self-supervised learning. b) The 3D U-Net is then trained to recover the manually masked region by learning the underlying sine wave patterns. A combination of loss functions is employed: the Generative Adversarial Network (GAN) loss ensures the realism of the recovered features, while other losses are used to improve feature and style consistency across both sinogram and projection planes. c) Results of the recovered masked sinograms compared with the original ground truths. The model shows it can realistically reconstruct the masked sinogram through self-supervised learning. d) SXT Imaging and Tilt Series Coordinates. Soft X-ray tomography was performed by illuminating the specimen with a Soft X-ray light source and acquiring a tilt series of two-dimensional (2D) projections. The attainable imaging angular range is typically limited, resulting in missing wedges (red arrows). The three-dimensional (3D) tilt series stack was defined as follows: the Z-axis was positioned parallel to the optical axis and perpendicular to both the detector plane and the zero-tilt projection. The Y-axis was aligned parallel to the rotation axis, and the X-axis was defined as the horizontal axis perpendicular to both the Z- and Y-axes.

This work utilizes a 3D U-Net [54][55] architecture for the inpainting of the 3D tilt series sinogram. However, the standard implementation employing transposed convolutions in the decoder produced checkerboard artifacts in the output, as previously noted in the literature [56]. To mitigate these artifacts, the transposed convolution layers were replaced with the nearest neighbor interpolation as the upsampling method.

Four distinct loss functions were implemented during training (Fig. 1b). The Generative Adversarial Network (GAN) loss and the L1 loss are calculated directly between the output (inpainted 3D tilt series stack) and the target (original 3D tilt series stack). On the other hand, the perceptual loss and style loss [57] are derived from features extracted by a pretrained 2D VGG network [58]. To ensure feature consistency across different perspectives, the perceptual and style losses are computed from two perspectives: the sinogram perspective (XZ-plane, Fig. 1d) and the projection perspective (XY-plane, Fig. 1d).

### Sinogram Inpainting at Missing Angles

Next, we assessed the model’s ability to effectively recover randomly occluded sinogram sections (Fig. 1c). Despite the phase of the recovered sinogram being randomly selected, the model demonstrates its ability to consistently reconstruct the accurate sine-wave pattern on unseen test datasets. This outcome suggests that the model possesses a fundamental understanding of the sine-wave properties, rather than simply relying on rote memorization of the input data.

The core challenge for missing wedge correction through sinogram inpainting lies in generalizing the learned self-supervised features to the missing-angle sinogram sections (Fig. 2, red boxes of inpainted sinogram). Given the inherently limited angular range, the model lacks training exposure to the sinogram’s edge angles. Successful reconstruction, therefore, requires the model to extrapolate the discovered sine-wave patterns from the visible data to accurately infer the information within the missing wedge.

**Figure 2.**
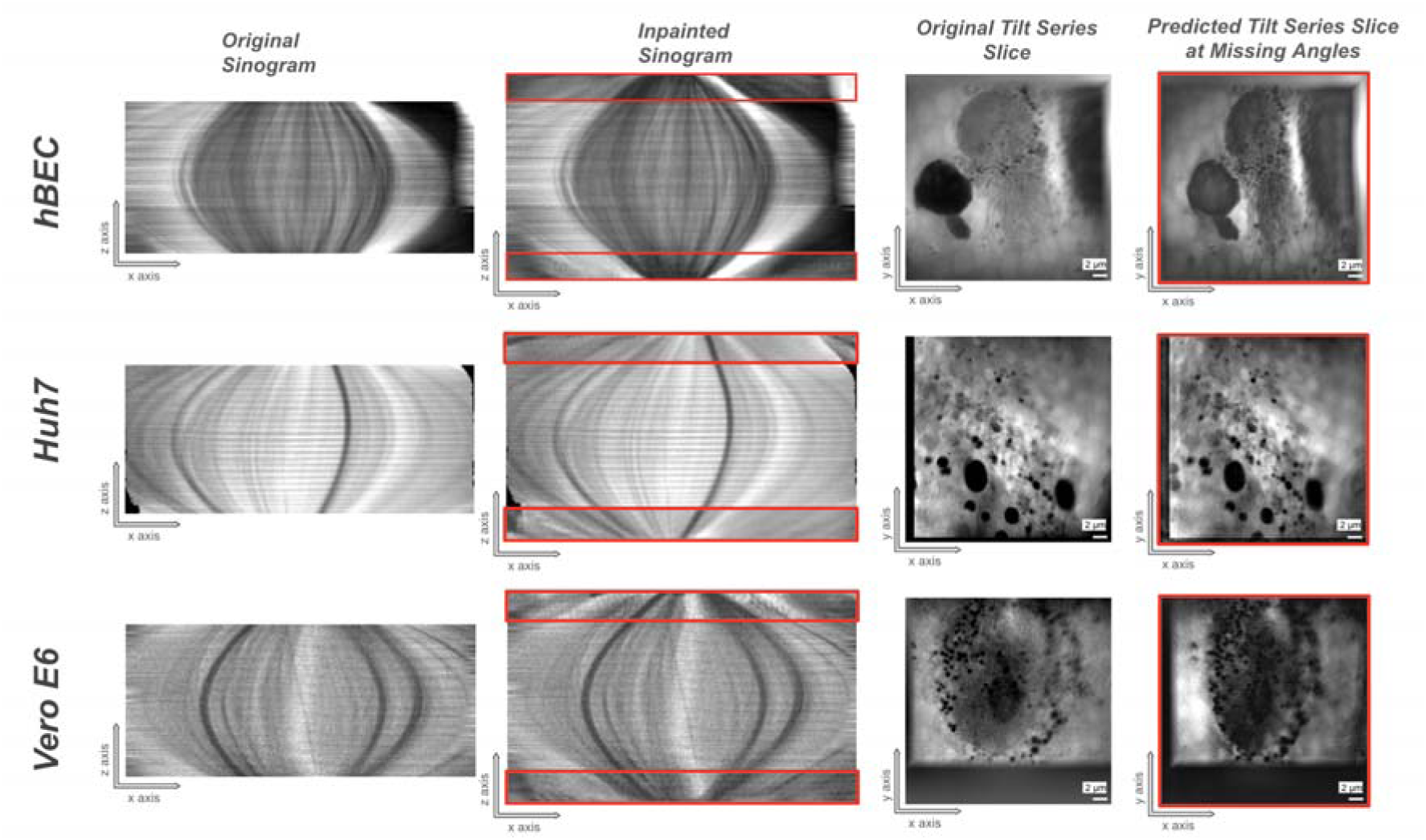
Sinogram Inpainting at Missing Angles on Diverse Cellular Structures (Cell Types from top: hBEC, Huh7, Vero E6). The column “Original Sinogram” shows the real sinogram with missing angles. The missing region sits on both sides of the sinogram (as shown in the red boxes in the “Inpainted Sinogram” column. The goal is to apply the model trained on known regions of the sinogram to the missing angles, with the knowledge of sine-wave patterns learned from self-supervised training. The column “Inpainted Sinogram” demonstrates our model’s generalizability to the missing angles and recovers natural sine-wave patterns following the original sinogram. The visualization of the XY-plane (columns “Original Tilt Series Slice” and “Predicted Tilt Series Slice at Missing Angles”) provides a more intuitive way of verifying the prediction of missing-angle slices. While following the perception angle shift, consistent cellular structures on various cell types are also maintained throughout the predicted missing-angle slices.

We applied the model trained on known regions to the missing-wedge region of several mammalian cell lines containing various internal membrane structures. Inpainted tilt series stacks are shown in Fig. 2 (inpainted sinogram). The model demonstrated convincing sine-wave patterns in the originally missing sinogram regions in the XZ plane of the tilt series stack (plane coordinates are defined in Fig. 1d). Furthermore, we visualized the XY-plane of the tilt series stacks, and observed the angle shifts on cellular landscapes to more intuitively verify the correctness of the predictions (Fig. 2, predicted tilt series slice at missing angles). It shows a natural perception angle shift following the original tilt series slices and maintains consistent cellular structures throughout the inpainted missing-angle slices on various cell types.

To further assess prediction accuracy, we analyzed the shape changes of lipid droplets (LDs) on tomograms reconstructed from the original, 40° inpainted, and 60° inpainted tilt series. LDs are ideal verification candidates because their known spherical morphology allows for quantitative measurement of distortion using metrics such as eccentricity and axis ratio. Furthermore, LDs are highly discernible in soft X-ray tomograms: their high carbon density results in a distinct, high X-ray absorption, making them the darkest features. This consistent contrast provides a characteristic Linear Absorption Coefficient (LAC) range, enabling straightforward segmentation via intensity thresholding [59].

We collected a total of nine tomograms of Huh7 cells exhibiting clear LDs. These tomograms were reconstructed from the original, 40°-inpainted, and 60°-inpainted tilt series. To quantify the missing-wedge distortion, we analyzed the LD shapes within the XZ-plane of the tomograms, as elongation is most pronounced along the optical axis (Z-axis), parallel to the zero-tilt projection direction. LD masks were derived by semi-automatic segmentation, employing Otsu’s method (detailed further in the Methodology section), yielding a series of 2D LD patches for shape analysis.

We calculated the eccentricity and axis ratio (minor axis / major axis) of the segmented LD masks. These two metrics served as quantitative indicators of elongation, based on the observation that the missing-wedge effect primarily distorts the spherical LDs by elongating them along the Z-axis in the XZ-plane. LDs reconstructed with the inpainted tilt series (40° and 60°) exhibited a statistically significant (Fig. 3a, b. Both with p values approaching 0 in eccentricity and axes ratio, compared to the LDs in the original tomograms in two-tailed paired t tests) improvement over the original missing-wedge series in both eccentricity and axis ratio (Fig. 3a, b). These results confirm that the sinogram inpainting successfully restores the LD morphology toward its native spherical shape.

**Figure 3.**
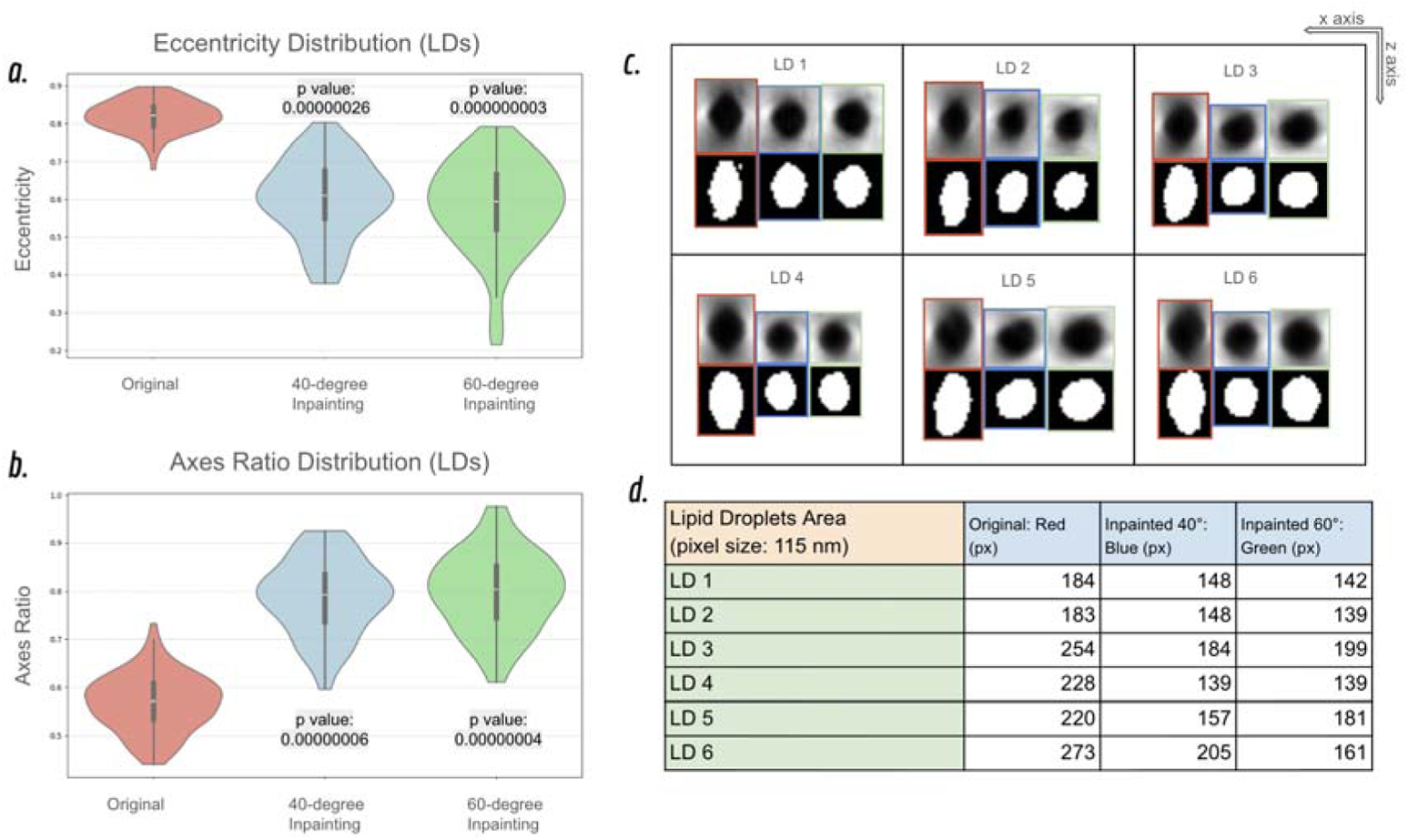
Restoration of Lipid Droplet (LD) morphology following Sinogram Inpainting from Huh7 cell. LDs reconstructed from tilt series that underwent sinogram inpainting were compared against the LDs with the original tilt series. **a)** LD Eccentricity Distribution. The distribution shows a significant shift toward lower eccentricity values (closer to the ideal spherical value of 0) in the inpainted tomograms. This reduction in eccentricity indicates that the inpainting effectively mitigates the shape elongation caused by the missing wedge effect, restoring the LDs to their native spherical morphology. **b)** LD Axis Ratio Distribution. The distribution of the major-to-minor axis ratio further confirms the improvement, showing a clear reduction in the ratio for LDs in the inpainted tomograms, thus demonstrating enhanced sphericity. **c, d)** Qualitative LD Shape and Volume Comparisons. Visual inspection of the tomographic slices (c and d) confirms the quantitative findings. Lipid droplets (LDs) in the original tomogram exhibit significant elongation along the Z-axis (a well-known consequence of the missing wedge), whereas the LDs reconstructed from the inpainted tilt series clearly demonstrate restored spherical morphologies.

Qualitative verification, shown by comparing selected LD patches (Fig. 3c, d), further supports these findings. The elongation effect on the original tomogram (Fig. 3c) not only distorts the spherical shape but also artificially enlarges the apparent LD volume (or area in 2D). Conversely, LDs in the inpainted tomograms restore the correct spherical morphology by effectively removing the elongated tails along the Z-axis.

### Applications on Diverse Cellular Structures

To test the generalizability of the sinogram inpainting model and inspect its impact on various organelles, we applied it to tomograms of cells with distinctive internal structures. Figure 4 displays four such tomograms, each reconstructed from the original, 40°-inpainted, and 60°-inpainted tilt series. The tomograms presented here were initially imaged using a ±55° tilt range, resulting in 111 tilt series slices. After application of the two inpainting models (40° and 60°), the resultant tilt series contain 151 and 171 slices, respectively. While SXT imaging angles can theoretically reach ±70°, the actual attainable angle is dependent on the sample grid position and sample holder design. For experimental convenience and practicality, we utilized the ±55° imaging angles, which still allow our model to demonstrate its effectiveness, rather than pursuing the maximum possible angles to achieve a full-angle inpainting.

**Figure 4.**
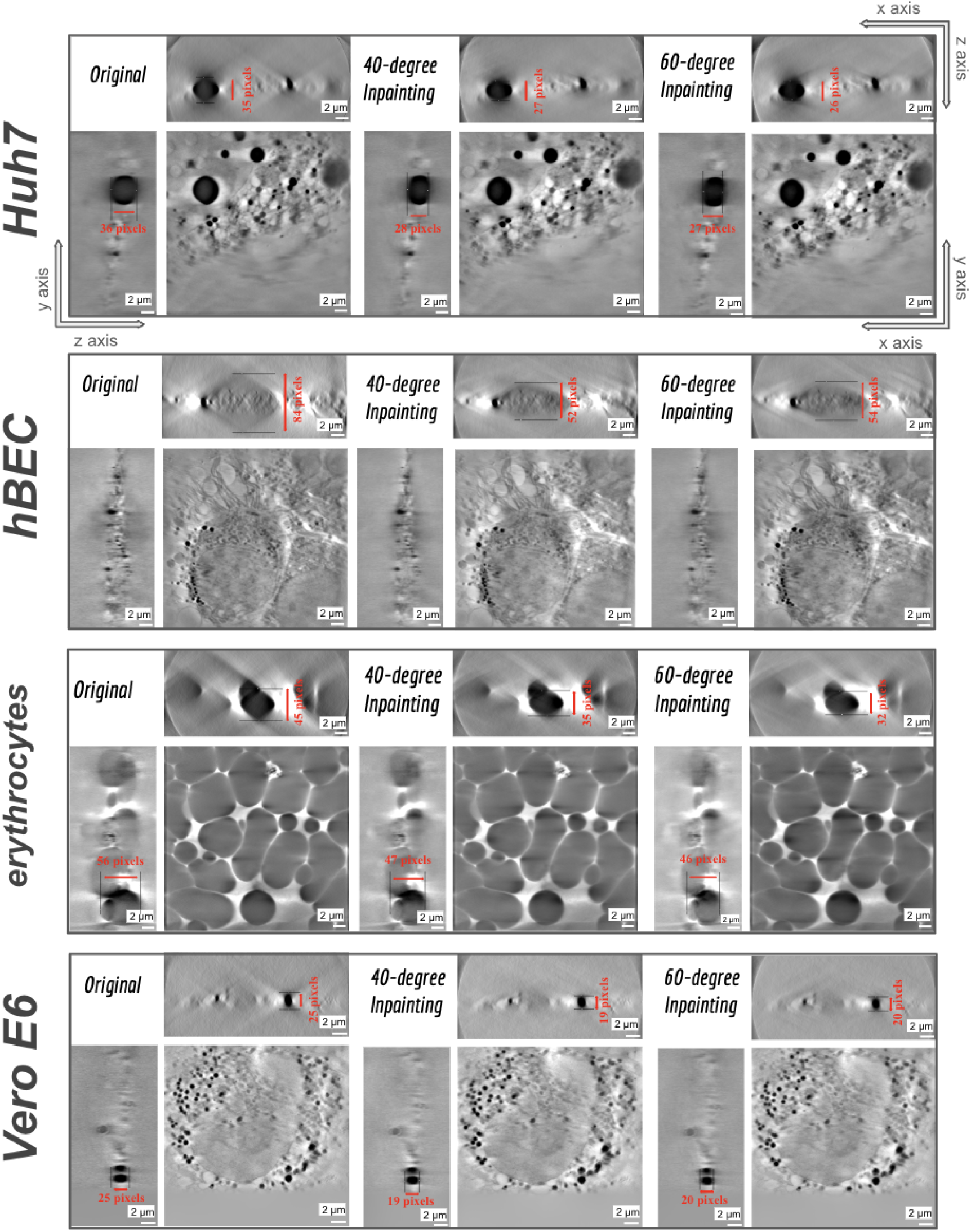
Mitigation of the Missing Wedge Effect on Various Cellular Structures (Cell Types from top: Huh7, hBEC, erythrocytes, Vero E6). Tomograms reconstructed from the original, 40°-inpainted, and 60°-inpainted tilt series are visualized across three orthogonal planes to detail the impact of sinogram inpainting. The displayed perspectives are the XZ-plane, YZ-plane, and XY-plane (axis coordinates shown in the first row). The missing wedge effect causes cellular structures in the original tomograms to appear elongated along the Z-axis, a distortion most pronounced in the XZ-and YZ-planes. Quantitative measurements of organelle lengths along the Z-axis demonstrate that the application of our sinogram inpainting model significantly reduces this elongation, effectively mitigating the missing wedge distortion.

To inspect the inpainting effect in detail, we visualize the tomograms across three orthogonal planes: the XZ-plane (top), the YZ-plane (left), and the XY-plane (providing a cellular structures perspective). Since the missing-wedge effect causes elongation along the Z-axis, its consequences are most pronounced and visually distinct in the XZ- and YZ-planes. We quantified this distortion by measuring the difference in organelle length along the Z-axis under the three conditions (original, 40° inpainting, and 60° inpainting) using various cellular features (Fig. 4). The results demonstrate that the inpainted tomograms significantly reduce the organelle length along the Z-axis. This evidence indicates that our sinogram inpainting method mitigates the missing-wedge effect that artificially enlarges organelle volumes, thereby restoring organelle shapes closer to their true morphologies.

We further applied our model to investigate its impact on measurements of *Plasmodium falciparum* hemozoin crystal volume. After invading the erythrocyte, *Plasmodium* digests up to 75% of the host hemoglobin to obtain essential amino acids for growth and protein synthesis [60]. Since the by-product free heme (Fe - protoporphyrin IX) is toxic to parasites, they detoxify it by transforming it into hemozoin crystals [8][9][10][11][12]. There is strong evidence that quinoline-based antimalarial drugs cap the surface of hemozoin crystals, inhibiting their growth, and causing the death of malaria parasites due to accumulation of heme and starvation from inability to catabolize hemoglobin [11][12][13][14][15][16][17][18][19]. Volumetric imaging of hemozoin crystals is an indicator of quinoline-type antimalarial drug activity through inhibition of their nucleation and growth.

While SXT is well-suited for this measurement, the missing wedge effect can introduce distortion into the reconstructed volume and subsequently bias quantitative analysis. This missing-wedge effect causes the crystals to appear elongated along the Z-axis in 3D reconstructions, leading to an overestimation of the true hemozoin volume, which can skew drug effect quantification.

In Figure 5, we present a detailed comparison of the missing wedge elongation effect across the original, 40°-inpainted, and 60°-inpainted tomograms. Visual inspection (Fig. 5, red text annotations) demonstrates that the elongation along the Z-axis is significantly mitigated by our sinogram inpainting model. To accurately quantify this effect across the entire crystal, we performed 3D segmentation of the crystals (Fig. 5b) using Otsu’s method [61] as described in the Methodology section. The 3D analysis shows that in the original tomogram, the crystal length is stretched along the Z-axis, leading to an overestimation of its true length.

**Figure 5.**
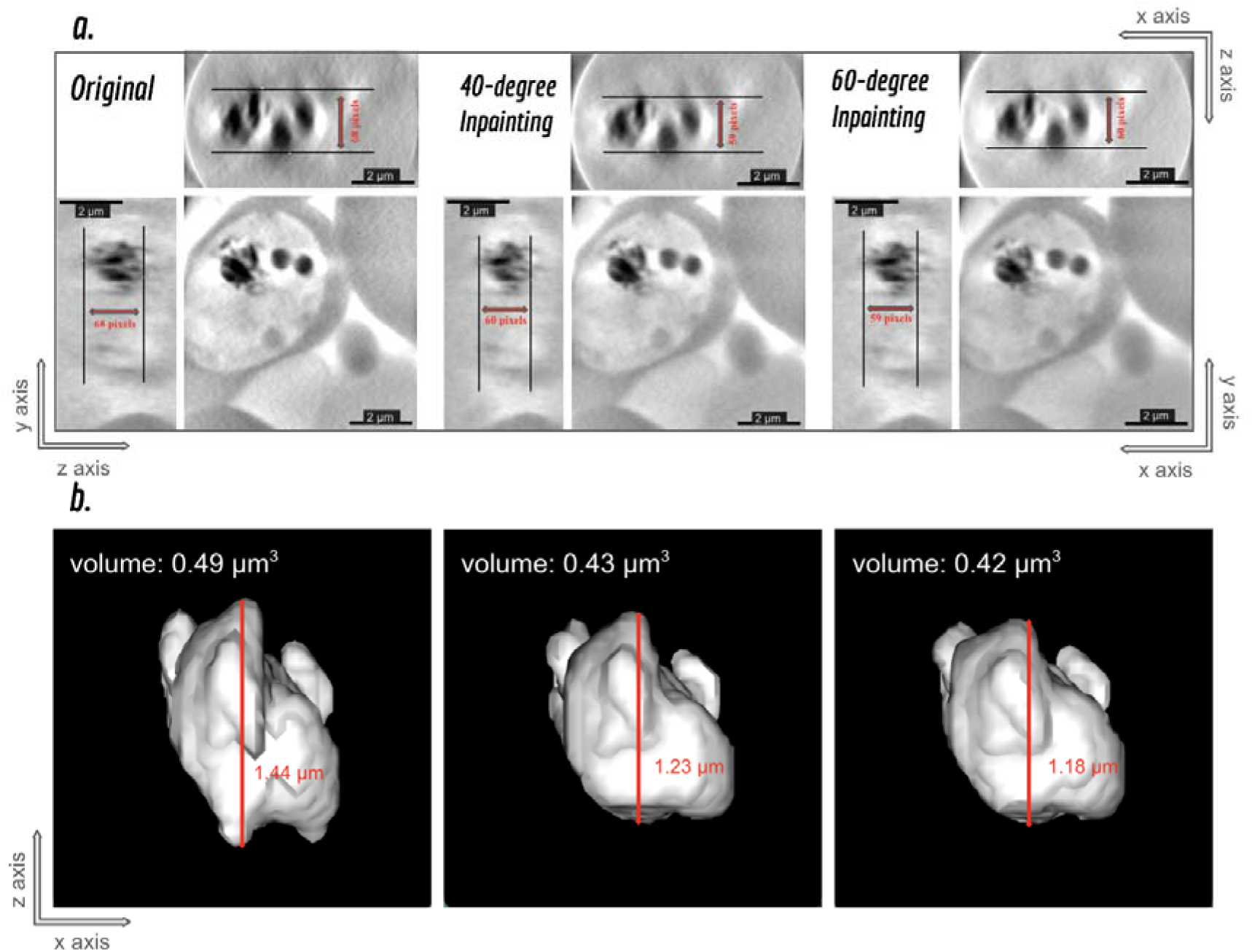
Mitigation of Missing Wedge Elongation in Hemozoin Crystals. Comparison of hemozoin crystals reconstructed from the original, 40°-inpainted, and 60°-inpainted tilt series, demonstrating the correction of morphological artifacts. **a)** Visualization across the XZ-, YZ-, and XY-planes shows that the elongation along the Z-axis is significant in the original tomogram. Both the 40°-inpainted and 60°-inpainted tomograms exhibit mitigated elongation, confirming the restoration of the crystal shapes toward their true morphologies. **b)** Quantification of Hemozoin Crystals with and without missing wedge correction. The difference in the crystal volume with and without the in-painting could be up to (0.49-0.42)/0.42*100=16.6%, which indicates the importance of missing wedge correction in SXT.

Conversely, the 40°- and 60°-inpainted tomograms show restored crystal structures, reversing the elongation artifact present in the original tomogram and providing a more accurate representation of the true crystal length (Fig. 5b). Calculation of the volume of the segmented crystals shows that the increase in volume caused by the missing wedge effect could be up to 16.6%, which could impact the accurate observations of hemozoin crystal growth rate.

## Discussion

In this work, we propose a novel sinogram inpainting framework for SXT designed to mitigate the missing wedge elongation artifact inherent in reconstructions of highly heterogeneous cellular structures. This mitigation also enables more accurate morphological and volumetric quantification. We demonstrate two models that increase 40°and 60° tilt series angle, respectively, both showing significant improvements with the missing-wedge elongations. In practical SXT imaging, the attainable imaging angle range depends on many factors, such as grid position and sample holder design. Therefore, the current model version (40°and 60°) might not be able to reach full-angle inpainting in all scenarios.

However, even if not reaching full angle, our models still show significant improvements from the original tomograms. Furthermore, the same framework can be adjusted to accommodate different angular ranges, depending on the imaging practices in specific scenarios and on the specific soft X-ray microscopes.

Although a direct ground truth sinogram for the missing-angle regions is unavailable to confirm the prediction, we demonstrate the reliability of our framework through two independent validation approaches: known region inpainting and lipid droplet shape restoration. First, the models demonstrate their ability to accurately recover masked regions within the original sinogram, confirming their correct understanding of the underlying sine-wave projection patterns. Second, and more critically, the models successfully restore the known spherical morphology of lipid droplets (Fig. 3), which serves as a compelling internal standard to further confirm their reliability in mitigating the missing-wedge effect. Lastly, we apply our sinogram inpainting model to various distinctive cellular structures (Fig. 4), and by measuring the cellular length along the Z-axis, we observed that our missing wedge correction models significantly mitigate the known elongation effect that previously overestimates the cellular length and volume. Furthermore, by applying our model to reduce the missing wedge effect in soft X-ray tomograms of *P. falciparum*, we corrected a 16% overestimation of hemozoin content caused by distortion in the reconstructed volume. This will allow a more accurate assessment of hemozoin crystal growth after being treated with antimalarial drugs. In future work, we plan to investigate whether chloroquine, a quinoline-based antimalarial drug, prevents or significantly slows down hemozoin crystal growth in *P. falciparum*, and to what extent [11][12][13][14][15][16][17][18][19]. The missing wedge correction approach will enable us to quantify changes in hemozoin crystal volumes below the uncertainty of ca. 16%, which results from the missing wedge artifacts. This is important because *P. falciparum* consumes hemoglobin and produces hemozoin crystals over an extended period (ca. 15h) as a means of heme detoxification.

In conclusion, the proposed sinogram inpainting framework represents an advance in SXT imaging and analysis. By effectively mitigating the missing wedge elongation artifact, this method enables more accurate cellular morphology and volume quantification, which were previously compromised [20][21][22][23][24][25][26]. These enhanced features are crucial across numerous biological and biomedical research areas. For instance, the accurate restoration of volume is vital for quantitative studies, such as determining the mode of action of anti-malarial drugs by observing volume changes in hemozoin crystals (Fig. 5). More broadly, the framework provides the necessary precision for reliable phenotyping of organelles and quantitative analysis of cellular pathways, offering a powerful tool for studying disease mechanisms, drug efficacy, and cellular responses to stimuli in a robust 3D context.

## Methodology

### Model Architecture

We opted for a 3D U-Net architecture over a 2D approach (such as UsiNet) primarily due to the highly heterogeneous nature of cellular structures in SXT. In this domain, even minor reconstruction artifacts can drastically compromise resolution and lead to the loss of critical biological information. Although a 3D model is more computationally expensive during training, it provides consistent inter-slice contextual information across the entire tilt series stack. This ensures structural coherence and minimizes artifacts between adjacent projections (XY-plane slices), which is essential for accurate reconstruction. A major practical advantage of the 3D model is the minimization of inference time, as the entire 3D tilt series stack can be processed in a single forward pass, eliminating the need for iterative slice-by-slice input. Furthermore, our one-step training pipeline and the sinogram-cropping training scheme synergistically decrease the necessary input data size and enhance training efficiency.

Our 3D Unet encoding path comprises a series of 3D downsampling stages. Each stage consists of two sequential ConvBlocks followed by a max-pooling operation. Each standard convolution block consists of a 3D convolution layer with a 3⋅3⋅3 kernel, followed by 3D batch normalization, and activated by the Exponential Linear Unit (ELU) function. On the other hand, the decoding path reconstructs the output resolution. At each level, the feature maps are upsampled, followed by a 5⋅5⋅5 convolution. It is worth noting that instead of the standard 3D Unet’s transposed convolution upsampling, we use the nearest neighbor upsampling because of significant checkerboard artifacts with the transposed convolution in our experiments and in previous studies [56]. The upsampled feature maps are concatenated with the corresponding encoder feature maps containing high-resolution information via a skip connection. This combined tensor is then processed by two additional ConvBlocks.

Because we adopt a Generative Adversarial Networks (GAN) training scheme, the above 3D U-Net is used as our generator. The discriminator architecture is a simple 3D convolutional network composed of five convolutional layers. Each layer uses spectral normalization to regularize the Lipschitz constant of its weight matrices, followed by a Leaky ReLU activation function.

### Model Training Scheme

We adopted a GAN framework for training the sinogram inpainting model, augmented by three additional loss functions — L1 loss, style loss, and perceptual loss — to ensure feature and style consistency, particularly in high-frequency details. While both the L1 and GAN losses are directly applicable to 3D data, the style loss and perceptual loss are calculated based on features extracted from a pre-trained 2D VGGNet. Consequently, we apply these two losses across two distinct 2D planes of the 3D tilt series stack (Fig. 2). Calculation on the XZ-plane (the sinogram perspective) is essential for maintaining the consistency and style of the underlying sine-wave projection patterns. Simultaneously, calculating the losses on the XY-plane (the cellular structures perspective) is crucial for preserving the consistency and detail of the cellular morphology throughout the missing-angle slices, thereby ensuring the quality of the final tomographic reconstruction.

We set the weights for the four loss components based on empirical observation of their respective scale and contribution: L1 loss*1, Perceptual loss*1, GAN loss*1, and Style loss*250. While these weights provided effective convergence, further experimentation could be conducted to optimize the loss weighting scheme. The maximum learning rate was set to 0.001, and a linear learning rate schedule was applied across a total of 20,000 training epochs. Although 20,000 epochs were applied, we observed that the model produced realistic sinogram predictions at the missing angles after approximately 10,000 epochs, with subsequent training primarily refining fine-scale details. Importantly, no deterioration in performance was detected even after the full 20,000 epochs.

### Tilt Series Data Processing

Due to the significant computational cost associated with the 3D U-Net architecture, we experimented with several strategies to manage the input tilt series dimensions. Based on the results and practical usage, we proposed a scale-invariant training strategy that balances the trade-off among the desired field of view (FOV), the preserved resolution, and the potential for introducing inter-stack artifacts along the Y-axis.

Because of the bottleneck of GPU RAM, the spatial dimension of 3D input data (tilt series) is limited (256*256 or 512*512 in our experiments). Two options could process our original dimension (1024*1024) into this limited spatial size: interpolation (loss of resolution) and center-cropping (loss of FOV). However, we observed that from the perspective of the sinogram, the fundamental sine-wave patterns learned by the model remain identical regardless of whether the tilt series is interpolated or cropped to a smaller size. Therefore, we propose a scale-invariant training scheme: during training, we first interpolate the spatial dimension between the range (256 or 512, 1024) and then use a center-crop (256*256 or 512*512) to ensure a consistent input dimension. In this way, the model can perform sinogram inpainting across different scales in inference time, depending on the need for resolution or FOV. For instance, when prioritizing resolution, the input can be center-cropped to preserve the original resolution (such as in Fig. 5). On the other hand, when FOV is prioritized, the input can be interpolated at the cost of loss of resolution. Alternatively, both techniques (center-cropping and interpolation) can be used together to balance the loss of resolution and FOV, which is the major advantage of our scale-invariant training. An alternative option to maintain both original resolution and intact FOV is by dividing the tilt series stack into smaller sub-stacks along the Y-axis. While this maximizes both parameters, it introduces slight artifacts between sub-stacks and increases the overall training duration. Overall, we found that models trained on smaller images already offered substantial practical utility, providing flexible application modes that deliver complementary information essential for biological insight, while full-resolution and full-FOV models can also be achieved with longer training time.

### Segmentation of Lipid Droplets (LDs) and Hemozoin Crystals

Accurate measurements of LDs and hemozoin crystals are important for evaluating our model’s performance. While both objects appear darkest in SXT images, it is still difficult to use a simple threshold to cleanly segment them because there are other objects dense in carbon that also appear as dark objects, such as gold nanoparticles. In this work, we adopt a 2-step thresholding segmentation that best removes unwanted objects and only segments LDs and hemozoin crystals.

The segmentation of target objects (LDs or hemozoin crystals) was achieved through a multi-step thresholding approach designed for robust noise exclusion and precise object definition. Initial filtering involved applying a percentage-based intensity threshold, specifically discarding pixel values above the 97th to 99th percentile, followed by a morphological filter to remove small-volume noise (e.g., gold nanoparticles) by excluding objects with a major axis length of 5 to 10 pixels. For the remaining candidate objects, we used their bounding box coordinates to perform localized, adaptive segmentation via Otsu’s method [61]. This provided an objective, automatically selected threshold for the distinctive segmentation of each target. We note that minor variations in tomogram contrast, potentially arising from sinogram inpainting, necessitated slight, empirical adjustments to the initial percentage intensity and major axis length thresholds across different tomograms to ensure optimal noise suppression and object integrity.

### Soft X-Ray Tomography Imaging

The tomogram data were collected using SXT on the SiriusXT SXT-100 laboratory-based microscope. Tomograms were acquired from flat samples grown on transmission electron microscopy (TEM) grids by projecting soft X-rays through the samples across a broad tilt series, spanning from ca. −55° to +55° with 1° increment at each step. Total tomogram acquisition times varied between one and two hours.

### Huh-7 Cell Sample Preparation

Human hepatoma Huh-7 cells were cultured as described previously in Dulbecco’s modified Eagle’s medium (DMEM) [62][63], supplemented with HEPES, non-essential amino acids, 100 U/ml penicillin/streptomycin and 10% fetal calf serum (complete medium) in a humidified incubator at 37°C and 5% CO_2_ Huh-7 cells (3×10^4^ cells/well) were seeded onto gold Quantifoil R 2/2 holey carbon-film microscopy grids (Au-G200F1) using one grid per well in a 12-well plate. Grids were vitrified by plunge-freezing in liquid nitrogen-cooled liquid ethane using Leica GP2 freezer.

### HBEC Cell Sample Preparation

Primary human bronchial epithelial cells (hBECs; FC-0035, CellSystems) were cultured as previously described [64]. Briefly, cells were expanded in PneumaCult-Ex Plus basal medium supplemented with hydrocortisone and primocin. For differentiation, hBECs were seeded onto polyester membrane inserts. An air–liquid interface (ALI) was established by removing the apical medium and feeding cells exclusively from the basolateral side. Upon confluence, the medium was switched to PneumaCult-ALI maintenance medium supplemented with hydrocortisone, primocin, and heparin. Cells were fed basolaterally three times per week. After four weeks at ALI, successful differentiation and ciliation were confirmed by light microscopy. Mature cells were then trypsinised, pelleted, and resuspended in PBS containing protease inhibitors to a final concentration of 1,000–2,000 cells/µL. For sample preparation, 3 µL of the cell suspension was applied to Quantifoil EM grids (3.5/1), excess liquid was blotted, and grids were vitrified by plunge freezing using a Gatan system.

### Vero E6 Sample Preparation

The Vero E6 cell line was obtained from ATCC (American Type Culture Collection, Manassas, VA, USA) and was maintained in Dulbecco’s Modified Eagle’s Medium (DMEM) supplemented with 10% Fetal Bovine Serum (FBS) and 1% penicillin/streptomycin (P/S), in a humidified incubator at 37°C and 5% CO_2_. For sample preparation, cells were grown onto 3 mm TEM grids, preliminarily irradiated for 3h with UV inside a biocabinet for sterilisation. Cells were seeded onto the sterile grids in 6-well plates at a density equal to 2×10^5^ cells per well in DMEM supplemented with 10% FBS and 1% P/S. The 6-well plates with the samples were placed in the incubator for 24h at 37°C and 5% CO_2_ to let the cells adhere to the grids.

For cryo-preservation for SXT imaging, at the end of the incubation time, the grids, with the cells attached onto them, were picked up from the 6-well plates, washed once in room temperature phosphate-buffered saline (PBS) and vitrified immediately by plunge-freezing with Leica GP2 freezer (9s blotting, 95% sample chamber humidity).

### Malaria Parasite Culture

Long-term in vitro-adapted *P. falciparum* clone 3D7 was maintained in culture as described previously [66]. In brief, parasites were grown at 5% hematocrit in serum-free RPMI-1640 medium and maintained at 37°C under controlled atmospheric conditions (2% O_2_, 5% CO_2_, 93% N_2_). Synchronised late-stage infected erythrocytes, purified by magnetic columns [65] and adjusted to 18% parasitemia and 1% hematocrit, were mixed with serum-free RPMI-1640 medium. After 2h and 4h at 37°C, infected erythrocytes were collected, washed twice in phosphate-buffered saline (PBS), and the pellet was diluted to 50% hematocrit with PBS before freezing.

For cryo-preservation for SXT imaging, cell samples were applied onto 3 mm TEM grids, preliminarily treated by glow discharging (Ar atmosphere, 30 seconds, 10 mA), and vitrified immediately by plunge-freezing in Leica GP2 freezer (6s blotting, 80% sample chamber humidity).

## Data availability

The data supporting the findings of this study are available from SiriusXT, Dublin, Ireland; however, restrictions apply to their availability, as they were used under license for the current study and are not publicly accessible. Data are, however, available from the corresponding authors, ShaoSen Chueh and Sergey Kapishnikov, upon reasonable request and with permission of SiriusXT.

## Acknowledgements

SSC, JCS, and SK acknowledge funding from the European Union’s Horizon Europe Research and Innovation programme under the Marie Skłodowska-Curie Actions Doctoral Networks (MSCA-DN, agreement No. 101120151).

AZ acknowledges Enterprise Ireland for funding under the Innovation Partnership Programme (grant number IP20241077).

## Author Contributions

***SSC***: Conceptualization, Data Curation, Formal Analysis, Investigation, Methodology, Software, Writing – Original Draft Preparation.

***MLP***: Resources (biological samples), Writing – Review & Editing (including description of the biological sample preparation in the methodology section).

***CdC, TI, AZ, VC, GC, PG***: Resources (biological samples), description of the biological sample preparation in the methodology section.

***CM, DR, SO, MD***: Conducting soft X-ray imaging of all samples presented in this work.

***JCS***: Supervision, Writing – Review & Editing

***SK***: Conceptualization, Data Curation, Formal Analysis, Validation, Supervision, Project Administration, Writing – Review & Editing.

## Declarations

The authors declare no competing interests.

